# A Hybrid Diffusion Model for Stable, Affinity-Driven, Receptor-Aware Peptide Generation

**DOI:** 10.1101/2024.03.14.584934

**Authors:** R Vishva Saravanan, Soham Choudhuri, Bhaswar Ghosh

## Abstract

The convergence of biotechnology and artificial intelligence has the potential to transform drug development, especially in the field of therapeutic peptide design. Peptides are short chains of amino acids with diverse therapeutic applications that offer several advantages over small molecular drugs, such as targeted therapy and minimal side effects. However, limited oral bioavailability and enzymatic degradation have limited their effectiveness. With advances in deep learning techniques, innovative approaches to peptide design have become possible. In this work, we demonstrate HYDRA: a hybrid deep learning approach that leverages the distribution modeling capabilities of a diffusion model and combines it with a binding affinity maximization algorithm that can be used for *de novo* design of peptide binders given target receptors. As an application, we have used our approach to design therapeutic peptides targeting proteins expressed by Plasmodium falciparum Erythrocyte Membrane Protein 1 (PfEMP1) genes. The ability of our model to generate peptides conditioned on the target receptor’s binding sites makes it a promising approach for developing effective therapies for malaria and other diseases.

## Introduction

As researchers delve deeper into the world of biotechnology and artificial intelligence, they discover exciting new drug development possibilities^1,2^. One particularly promising avenue is using deep learning techniques in peptide design. Peptides are short chains of amino acids that play critical roles in the body’s biological processes. They have already been developed and used as therapeutic agents for various medical conditions, thanks to their specific interaction with biological targets^3^.

The benefits of using peptides as drugs are numerous. Compared to conventional small-molecule drugs, peptides have fewer side effects^4^. They can mimic or interfere with naturally occurring peptide hormones or proteins in the body, influencing many physiological processes. Plus, when peptides are broken down, they produce amino acids, the body’s natural building blocks, which are well-tolerated. Prior studies have already explored numerous therapeutic applications for peptides, including treating autoimmune illnesses, metabolic diseases, cancer, and infectious infections.

Peptide therapeutics have revolutionized the treatment of various conditions, such as diabetes and bacterial infections. Insulin, for example, has been a life-saving drug for patients with diabetes, while peptide-based antibiotics have effectively treated bacterial infections^5^. Glucagon-like peptide-1 (GLP-1) analogs have also been used to manage type 2 diabetes and obesity^6^ with great success. In addition to their current applications, peptides are being studied for their potential as vaccine ingredients^3^, which could greatly benefit public health. Thanks to the use of deep learning, protein engineering has seen tremendous advances, allowing researchers to design reliable sequence and structure prediction techniques. State-of-the-art fast structure prediction techniques such as AlphaFold2 have played a crucial role in this progress, granting greater access to protein structures and accelerating research involving the design of new proteins and peptides^7–12^.

Both sequence-based^13^ and structure-based protein design approaches have been explored in previous studies. PepMLM^13^, a target sequence-conditioned peptide binder generating model, utilizes the Masked Language Modeling strategy to position peptide sequences at the terminus of target protein sequences to reconstruct the binder. PepMLM can generate binders for target proteins without requiring its structure, facilitating programmable proteome editing applications. EvoDiff^8^ attempts to merge evolutionary-scale data with diffusion models to design proteins. It can generate diverse and structurally plausible proteins covering a wide range of natural sequences and functions^8^, as well as proteins with disordered regions and functional structural motifs, which traditional structure-based models struggle with. RFdiffusion^7^, a model designed to fine-tune the RoseTTAFold structure prediction network for protein structure denoising tasks, is the most promising development in this field at the time of writing. RFdiffusion generates protein backbones and performs well in a wide range of tasks such as unconditional and topology-constrained protein monomer design, protein binder design, symmetric oligomer design, enzyme active site scaffolding, and symmetric motif scaffolding for therapeutic and metal-binding protein design.

While existing approaches focus on being able to design whole proteins and enforcing structural constraints such as symmetry and functional motif encoding, therapeutic peptide design requires multiple other attributes to be taken under consideration. Peptides are composed of smaller amino acid chains than proteins but are larger than many small-molecule drugs. Because of their intermediate size, they can be more susceptible to protease degradation in biological fluids^14^. Additionally, unlike proteins, peptides have a linear structure that can make them more prone to degradation^15^. One important factor that determines their effectiveness as drugs is their half-life^14^, which refers to the time it takes for half of the peptide to be broken down or eliminated from the body. Peptides with longer half-lives can exhibit their therapeutic effects over a longer time, which can be especially important for medications that require prolonged activity to be effective. Peptides often contain a mix of hydrophilic and hydrophobic amino acids, which can affect their solubility and interaction with cell membranes^16^. This can impact their ability to pass through cell membranes and work effectively inside cells. Peptides with a slightly higher percentage of hydrophobic residues may have increased cell permeability^16^ and be more effective at interacting with cell membranes, as hydrophobic residues can interact very well with the hydrophobic regions of lipid bi-layers, enhancing the transit of the peptide across cell membranes^16^.

In response to the limitations of existing methods, we present HYDRA, a hybrid deep learning approach for *de novo* receptor-aware peptide design. HYDRA combines a diffusion model that generates target-aware amino acid residues with a binding affinity maximization step to ensure stable interactions between the target receptor and the generated peptide. Inspired by TargetDiff^17^, a target-aware molecule generation model, we seek to generate amino acid residues in continuous 3D space in a non-autoregressive and SE(3)-equivariant manner. Using Cartesian coordinates, we represent the target receptor as atoms and peptides as amino acid residue points in 3D space. We define a diffusion process for continuous residue coordinates and discrete residue types where noise is gradually added and learn the joint generative process with a SE(3)-equivariant graph neural network^17^. Once sampled, the generated amino acid residues are reassembled into the exact peptide structure via an iterative binding affinity maximization process. This two-stage, hybrid approach allows our model to produce high-quality, diverse, and stable peptide binders for target receptor proteins. An empirical evaluation conducted on the PepBDB dataset^18^ demonstrates that HYDRA surpasses baseline models in its ability to generate stable peptide binders with high binding affinities towards the target receptors.

To demonstrate the feasibility of our approach, we have used it to design several *de novo* peptide binders to engage Plasmodium falciparum erythrocyte membrane protein 1 (PfEMP1), which are majorly responsible for antigenic variations of malaria^19^. Malaria, predominantly transmitted through bites of infected female Anopheles mosquitoes, remains a significant global health burden, particularly in tropical and subtropical regions^20,21^. Its severity is underscored by its lethality, especially in young children and pregnant women. In 2019, the World Health Organization (WHO) reported an estimated 229 million malaria cases and 409,000 deaths, highlighting the urgent need for effective treatment alternatives^22^. Around 6.3 million cases were reported from the Southeast Asia region, majority of cases were present in India^23^. Furthermore, the emergence of parasite resistance to established antimalarial drugs like chloroquine and sulfadoxine-pyrimethamine poses a critical challenge. This resistance hampers treatment efficacy, potentially leading to prolonged illness, increased healthcare costs, and elevated mortality risks. Beyond immediate mortality, malaria can have lasting detrimental effects on individuals, even in non-fatal cases. Recurrent infections can contribute to anemia, cognitive decline (particularly in children), and other complications, ultimately diminishing quality of life and economic productivity^24^. Drug resistance presents a central obstacle in malaria control. The effectiveness of new drugs wanes as the Plasmodium parasite develops resistance mechanisms. We have generated peptide drugs based on the PfEMP1 target proteins using HYDRA.

## Results

HYDRA’s peptide design pipeline, as illustrated in Fig. 1, starts by identifying suitable binding pockets (strong or medium affinity) within the target receptor protein. The chosen pocket is then extracted and fed into the model. A diffusion model takes the extracted pocket as input and generates the 3D positions and features for each amino acid residue, specifically tailored to fit the pocket’s topology. Finally, the generated amino acids are assembled into the final peptide structure using a binding affinity optimization algorithm that connects residues while maximizing the magnitude of binding affinity between the resulting peptide and the target receptor pocket.

**Figure 1.**
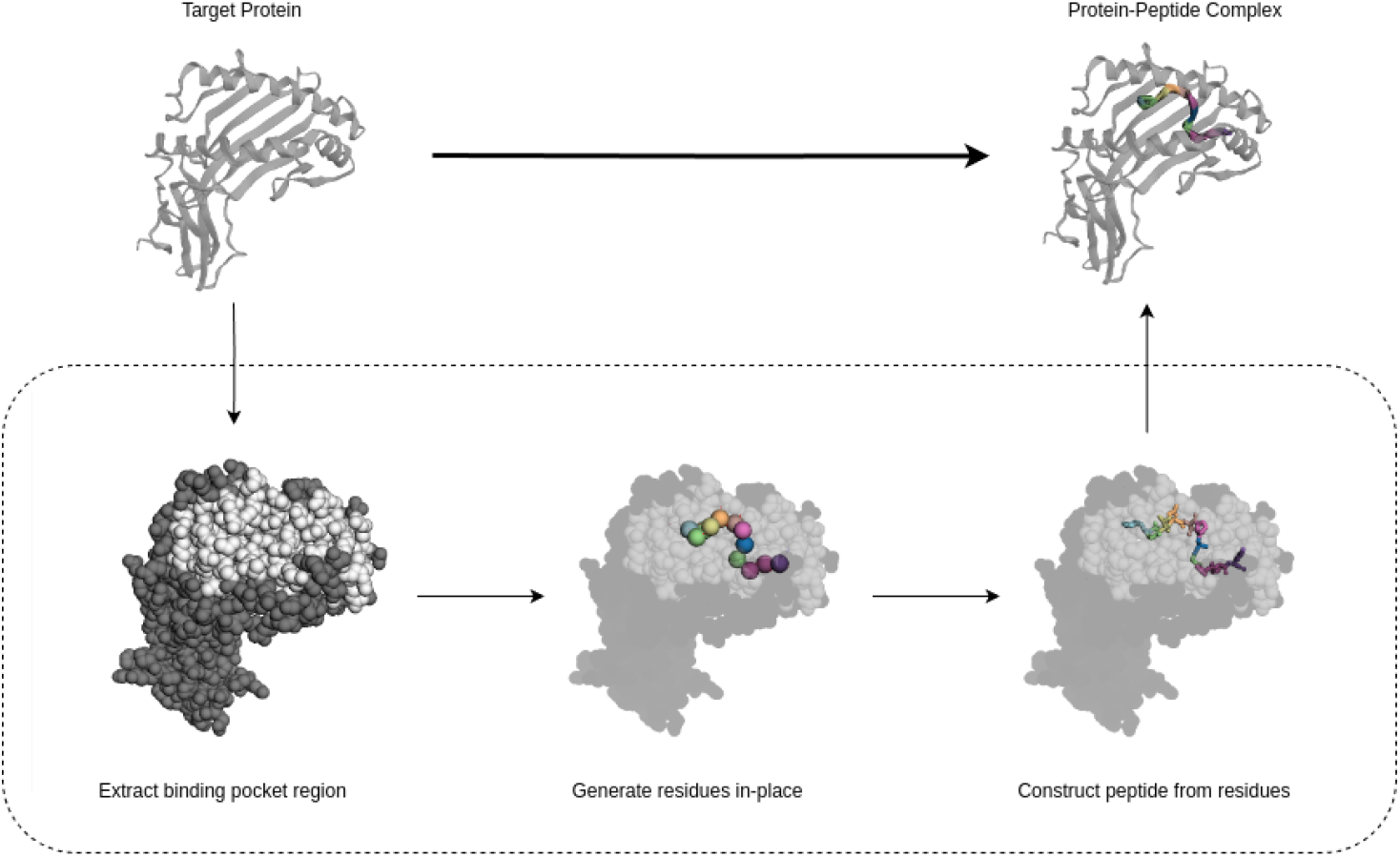
Overview of the peptide design process using HYDRA. The binding pocket region is extracted from the target receptor protein. This pocket then serves as input for a diffusion model, which generates individual amino acid residues in-place with the pocket. Finally, an optimization algorithm reconstructs the best peptide candidate structure based on its predicted binding affinity with the target receptor.

### Dataset

We constructed our training and evaluation sets based on PepBDB^18^, a comprehensive database of biological peptide-mediate complex structures with peptide lengths of up to 50 residues^18^. We curated 9225 protein-peptide complexes for the training dataset, 200 complexes for the validation dataset as well as 193 complexes for the test dataset. Table 1 summarizes the analysis of various physicochemical properties of the dataset’s peptides. A detailed description of the data curation methods is provided in the Supplementary Materials.

**Table 1.** Summary of different physicochemical properties of peptides in the PepBDB dataset.

| | MW [Daltons] | pI | $t_{1/2}$ [hours] | II | AI | GRAVY |
| --- | --- | --- | --- | --- | --- | --- |
| Avg. | 1337.39 | 7.13 | 10.12 | 48.88 | 81.63 | -0.41 |
| Med. | 1191.52 | 6.18 | 1.90 | 41.59 | 78.00 | -0.46 |

#### *In Silico* Assessment Criteria for Designed Peptides

We conducted a comprehensive computational evaluation of the generated peptides. This evaluation aimed to assess their potential as functional drug molecules in various applications. We employed various computational tools to analyze crucial physicochemical properties and binding affinities relevant to their potential therapeutic efficacy.

### Physicochemical Properties

#### Molecular Weight (MW)

The molecular weight of each peptide was calculated and compared to the established range (500-5000 Da) characteristic of drug-like peptides^25^. This parameter significantly influences a peptide’s solubility, membrane permeability, and potential for toxicity. Peptides falling outside this range might exhibit undesirable pharmacological properties, hindering their potential as viable drug candidates.

#### Isoelectric Point (pI)

The pI of each peptide was determined, reflecting the specific pH at which the molecule possesses a net neutral charge. This property is crucial in determining a peptide’s solubility, stability, and interactions with biological targets. Peptides with pI values outside the physiological pH range (7.4) might exhibit reduced solubility and stability in biological environments, compromising their therapeutic efficacy^26^.

#### Half-life (*t*_1/2_)

The half-life of a drug is the duration required for a drug’s concentration in the bloodstream (or any other pertinent compartment) to drop by half. The half-life of each peptide was calculated with respect to in-vitro conditions in mammalian reticulocytes. This parameter is critical for determining the efficacy and duration of action of a potential drug. Peptides with shorter half-lives might require more frequent administration to maintain their therapeutic effect, whereas those with longer half-lives could potentially offer sustained drug action, reducing dosing frequency and improving patient compliance. Most peptides have *in vivo* half-lives of 2–30 minutes due to protease enzymatic breakdown and quick renal clearance (molecules smaller than 30 kDa are quickly eliminated by glomerular filtration)^27^. Consequently, increasing the *in vivo* half-life of peptides to fulfill their therapeutic potential without requiring high dosages or frequent administration is desirable^27^.

#### Instability Index (II)

The instability index, based solely on the amino acid sequence of each peptide, was calculated using the formula 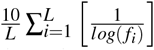, where L is the length of the protein sequence and *f*_*i*_ is the dipeptide frequency of occurrence for each dipeptide in the protein sequence. A peptide’s instability index is a numerical value that indicates how stable a protein or peptide will be^28^. Proteins or peptides are generally classified as stable if their instability index is lower than 40 and unstable if it is 40 or above^28^. Stable peptides are generally preferred for drug development due to their longer shelf life and potential for sustained activity *in vivo*. Peptides with high instability indices might undergo rapid degradation, limiting their therapeutic potential.

#### Aliphatic Index (AI)

The aliphatic index, reflecting a peptide’s overall hydrophobicity, is determined by calculating the proportionate volume that aliphatic side chains (Leucine, Isoleucine, Valine, and Alanine) occupy in a protein or peptide^29^. Due to the presence of hydrophobic interactions, peptides with higher AI may display improved structural stability, contributing to their overall stability, which is crucial for a therapeutic peptide to remain active. Peptides with higher AI values might also have higher membrane permeability, enabling them to reach intracellular targets effectively^29^.

#### Grand Average of Hydropathy (GRAVY)

GRAVY is a numeric representation of a protein or peptide sequence’s total hydrophobicity or hydrophilicity. This value is determined by the sum of the hydropathy of all amino acids divided by the total number of amino acids^30^. This value can be positive, negative, or zero. Positive GRAVY values indicate a hydrophobic sequence, whereas negative values show a hydrophilic sequence. A GRAVY near zero indicates a sequence with a balance of hydrophobic and hydrophilic residues. Positive GRAVY values indicate the sequence is dominated by hydrophobic residues, which may reduce the peptide’s solubility in aqueous solutions. Particularly at high concentrations, hydrophobic residues tend to group to minimize interaction with water molecules, which can cause protein aggregation. Hydrophobic residues can interact very well with the hydrophobic part of lipid bilayers, enhancing the transit of the peptides across cell membranes, which may have increased cell permeability. Extremely high hydrophobicity, however, may result in non-specific interactions with cell membranes that compromise the integrity of the membrane and impair cellular processes^30^. Negative GRAVY values indicate higher solubility in aqueous solutions because of the greater interactions with water molecules.

### Binding Affinity

Peptides exhibiting high binding affinity towards their target are more likely to disrupt crucial disease-associated processes and achieve therapeutic outcomes successfully. We assessed the binding affinity of each peptide towards its intended target using AutoDock Vina^31^ and FRODOCK^32^.

### AutoDock Vina

AutoDock Vina is a widely used program for molecular docking^31^. It employs an empirical scoring function to estimate binding affinity, considering factors like hydrophobic interactions, hydrogen bonding, and electrostatics. Lower scores (kcal/mol) indicate stronger predicted binding. Vina then performs local structure optimization using the Steepest Descent algorithm to refine the ligand’s pose within the binding pocket, seeking the pose with the lowest binding energy. While these scores are estimates, Vina provides a valuable tool for exploring potential ligand-receptor interactions^31^.

### FRODOCK

FRODOCK, or Fast Rotational DOCKing, is an approach for protein-protein docking simulations^32^. Unlike AutoDock Vina, which uses an empirical scoring function, FRODOCK employs a correlation function for protein-protein docking. This function assesses the overlap of interaction patterns (e.g., hydrophobicity, electrostatics) between protein surfaces, with higher scores indicating greater potential for favorable interactions but not directly reflecting binding affinity. This distinction highlights the importance of selecting the appropriate docking tool based on the specific type of molecular interaction under investigation^32^.

Utilizing two distinct scoring functions helps mitigate potential biases inherent to any single method and provides a more robust evaluation of predicted binding affinity.

#### Diversity

We quantify the diversity of the generated peptide sets using the average pairwise Tanimoto distance^33,34^. This metric assesses the similarity between peptide pairs by considering the presence or absence of specific amino acids at each sequence position. Higher average Tanimoto distances indicate a more diverse peptide set capable of exploring a wider range of potential binding interactions.

### Designing Peptides for Receptors in the Test Set

To assess HYDRA’s ability to generalize to unseen receptors, we employed a rigorous strategy. We focused on *de novo* peptide binder generation (novel peptides not encountered during training) for each of the 193 binding pockets within the independent test set. For each pocket, we generated 30 unique peptides, resulting in a total of 5790 novel peptides to evaluate HYDRA’s binding prediction for unseen receptors.

### HYDRA Outperforms RFDiffusion in Stable Peptide Design

To evaluate HYDRA’s performance relative to the current state-of-the-art, we compared it against RFDiffusion^7^. RFDiffusion leverages a diffusion model framework to iteratively refine candidate peptides for target receptor binding. It begins with a pool of random sequences and progressively adjusts them based on predicted binding scores, aiming to converge toward high-affinity binders over multiple iterations^7^. While RFDiffusion stopped at predicting just the peptide backbone, we went a step further by utilizing ProteinMPNN^35^ to generate full peptide sequences from these backbones for analysis of physicochemical properties.

Our evaluation revealed that peptides designed by HYDRA displayed significantly better binding affinities to receptor pockets compared to those generated by RFDiffusion. Additionally, HYDRA exhibited a slight edge in generating more diverse peptide sequences for each pocket. Analysis of physicochemical properties revealed that both models produced peptides within the established range for drug-like molecules in terms of molecular weight. However, HYDRA-designed peptides had isoelectric points closer to 7, indicating potentially greater solubility and stability compared to RFDiffusion-generated ones. Interestingly, the half-life data for both models showed a right-skewed distribution, meaning a few outliers skewed the mean values. While RFDiffusion peptides had a higher average half-life, HYDRA peptides boasted a higher median half-life, suggesting a more consistent level of stability across the generated set. Furthermore, HYDRA demonstrated a clear advantage in peptide stability. The percentage of stable peptides generated by HYDRA was significantly higher (58.31%) compared to RFDiffusion (31.02%). This trend was further confirmed by the lower instability index distribution for HYDRA peptides. Aliphatic Index values were comparable for both sets, with HYDRA having a slight edge in mean AI and RFDiffusion edging out on the median value. Finally, the GRAVY scores (generally negative for peptides) indicated higher aqueous solubility for RFDiffusion peptides compared to those generated by HYDRA.

A comprehensive comparison of both methods across various metrics is presented in Tables 2 and 3. Figures 2 and 3 depict the distribution plots for the aforementioned properties.

**Table 2.** Comparison of metrics for *de novo* peptides generated by HYDRA and RFDiffusion.

| Model | Vina Score (↓) |  | FRODOCK Score (↑) |  | Diversity (↑) |  | Stability (↑) |
| --- | --- | --- | --- | --- | --- | --- | --- |
|  | Avg. | Med. | Avg. | Med. | Avg. | Med. |  |
| HYDRA | **-4.112** | **-3.920** | **2277.171** | **2217.161** | **0.625** | **0.623** | **58.31%** |
| RFDiffusion | -2.764 | -2.745 | 1728.622 | 1772.440 | 0.579 | 0.600 | 31.02% |

**Table 3.** Comparison of different physicochemical properties of peptides generated by HYDRA and RFdiffusion.

| Model | MW [Daltons] | | pI | | $t_{1/2}$ [hours] (↑) | | II (↓) | | AI (↑) | | GRAVY | |
| --- | --- | --- | --- | --- | --- | --- | --- | --- | --- | --- | --- | --- |
|  | Avg. | Med. | Avg. | Med. | Avg. | Med. | Avg. | Med. | Avg. | Med. | Avg. | Med. |
| HYDRA | 1109.64 | 1099.28 | **6.97** | **6.00** | 10.89 | **3.50** | **41.35** | **35.09** | **90.39** | 80.00 | -0.20 | -0.21 |
| RFDiffusion | 1326.58 | 1258.94 | 6.07 | 5.49 | **17.65** | 1.90 | 84.88 | 73.58 | 88.55 | **86.67** | -0.93 | -0.86 |

**Figure 2.**
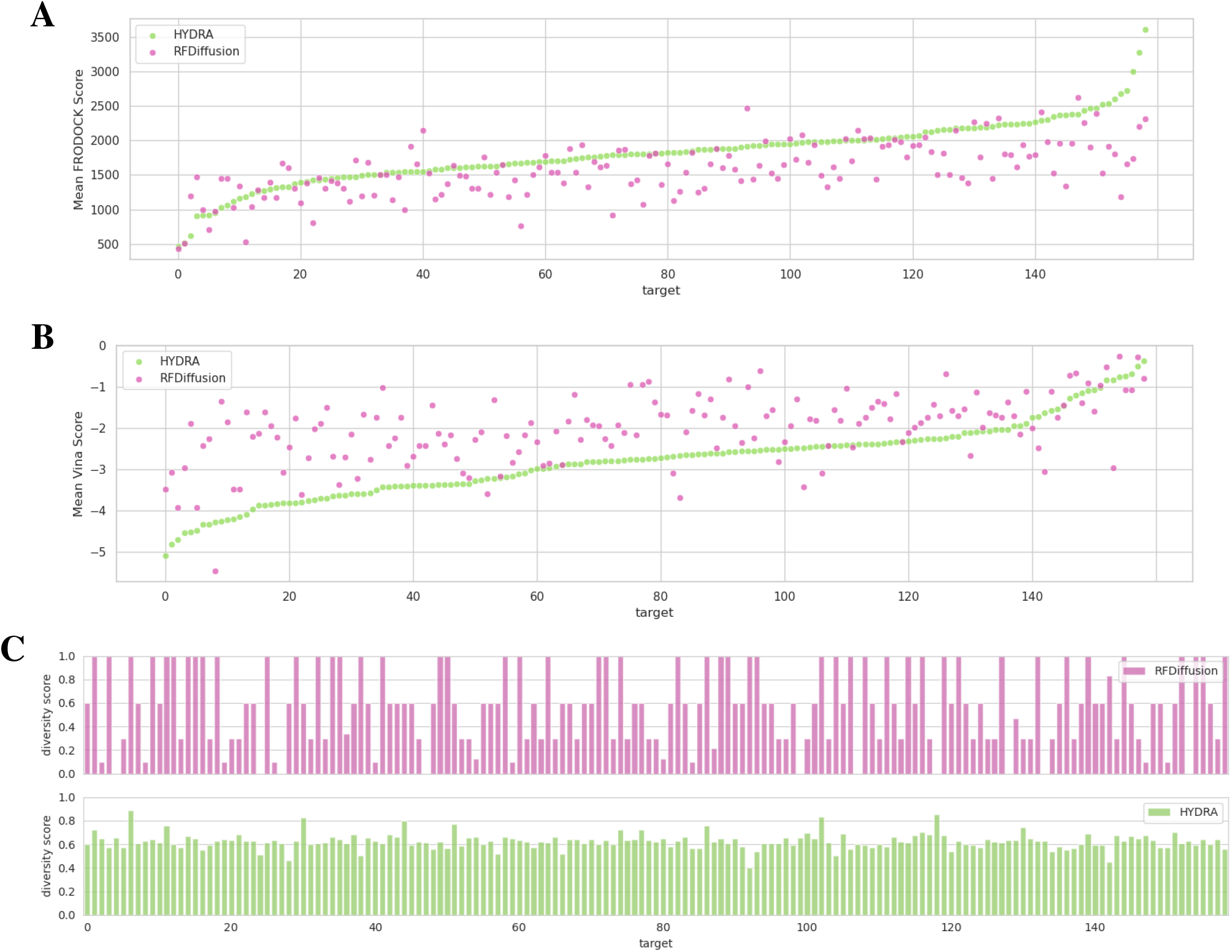
Comparison of binding affinity and peptide diversity between HYDRA and RFDiffusion. (*A*) and (*B*) Mean Vina and FRODOCK scores respectively for peptides generated by each model. Lower Vina scores and higher FRODOCK scores indicate stronger predicted binding affinities. Binding targets are sorted based on the mean score for HYDRA-generated peptides. (*C*) The pairwise Tanimoto diversity scores across the generated peptide sets for each binding target. While the average diversity scores are similar, the distribution of Tanimoto scores reveals greater consistency in HYDRA’s peptide diversity compared to RFDiffusion.

**Figure 3.**
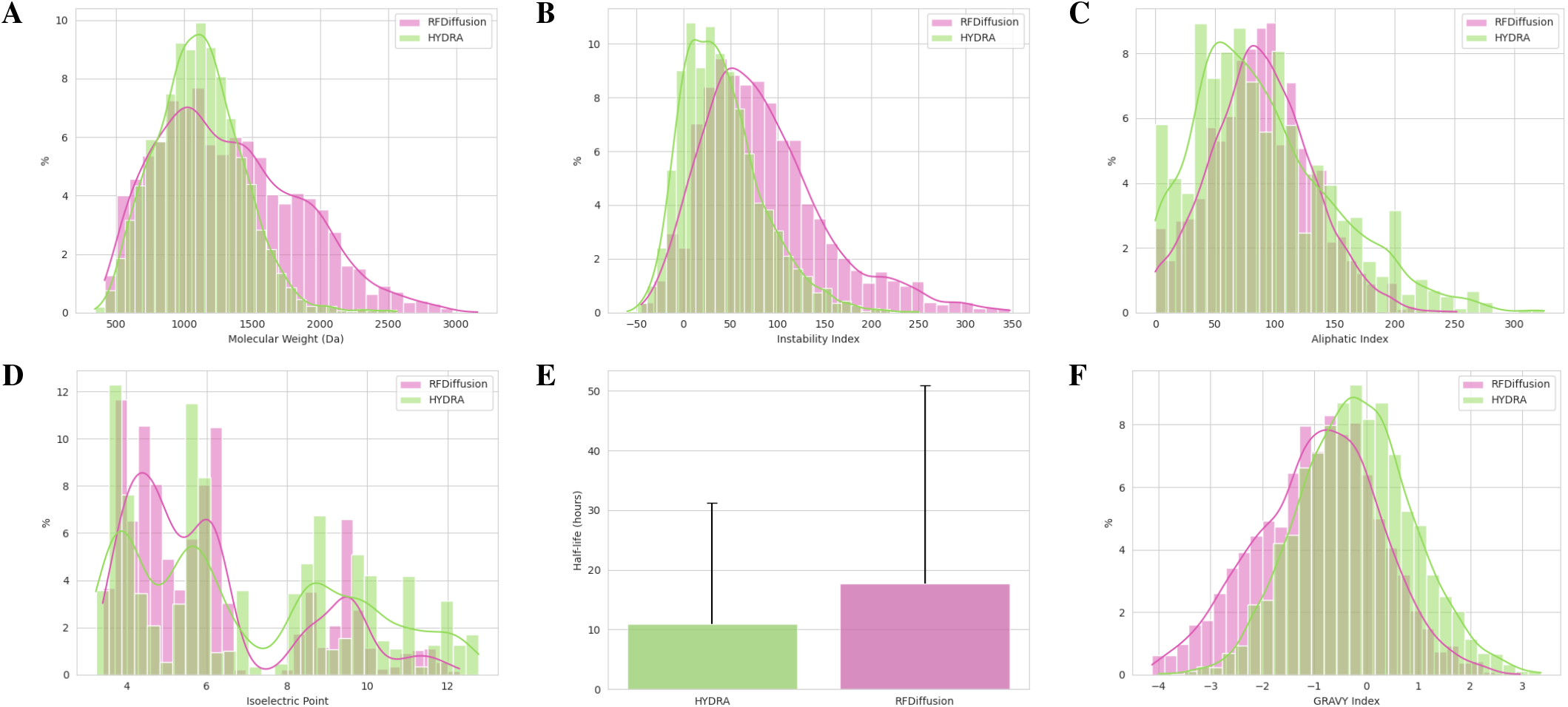
Comparison of physicochemical properties of peptides generated by HYDRA and RFDiffusion. Each histogram represents the distribution of peptides across different ranges for a specific property. The y-axis indicates the percentage of peptides within each range. (*A*) Molecular Weight in Daltons, (*B*) Instability Index, (*C*) Aliphatic Index, (*D*) Isoelectric Point, (*E*) Half-life in hours, (*F*) GRAVY Index. Peptides designed by HYDRA show a more favorable bias in their property distribution compared to RFDiffusion.

### Peptide Generation for PfEMP1 Targets

The malaria parasite, Plasmodium falciparum expresses a diverse family of PfEMP1 proteins with crucial roles in severe malaria pathogenesis. These 60 highly polymorphic surface antigens mediate cytoadherence of infected red blood cells to endothelial receptors, leading to sequestration and tissue damage^36^. We analyzed single-cell transcriptomics data of P. falciparum 3D7 to identify highly expressed PfEMP1s (PF3D7_1200600, PF3D7_1150400, PF3D7_70712400) based on cellular heterogeneity, cell-to-cell variability, and cellular states^37^. Using CAVITY^38^, we predicted strong and medium binding sites based on their druggability score (a higher score indicating stronger binding). We focused on these proteins due to available 3D structures from X-ray diffraction in the Protein Data Bank (PDB)^39^. Combining data from CAVITY and PDB, we identified extracellular domain binding sites crucial for potential drug interaction.

After identifying binding sites, our model generated candidate peptides. ExPASy^40^ then assessed their properties (molecular weight, isoelectric point, half-life, stability, and hydrophobicity). We prioritized stable peptides with high predicted binding affinities and favorable biophysical properties. Finally, to gain a complete understanding of their potential as therapeutic agents, we selected a few peptides per protein for further analysis. This multi-pronged pipeline included structural analysis, toxicity and immunogenicity assessment, and comparison with natural peptides.

Through this comprehensive selection process, we identified a final set of 12 candidate peptides (Table 4). These peptides are promising therapeutic agents targeting 3 distinct PfEMP1 variants. Furthermore, to assess the potential for broad-spectrum activity, we evaluated the binding affinity of these peptides with additional target receptors beyond those initially considered during their design. This analysis aimed to determine if any individual peptide could potentially bind to and interact with multiple binding sites, offering a broader therapeutic scope. Moving forward, the next crucial step in this research involves sending these selected peptides for wet lab experiments. This practical evaluation will be instrumental in validating our *in silico* findings and assessing the efficacy and safety of these peptides in a biological context.

**Table 4.** Identified binding site domains and target peptides selected following the filtering process.

| Target Protein | Binding Site Domain | Generated Peptide Sequences |
| --- | --- | --- |
| PF3D7_1200600 | Extracellular | SLGEVGAPALNFDA<br>ADPTPAHKEGS<br>LVVNTFVAGG<br>MDLTNPGANP<br>GPKTGGNGKSG |
| PF3D7_1150400 | Extracellular | GPMSAGAGATGM<br>NNGAGGDKTV<br>LGSTGMMVNP<br>GGSKFTGASTS |
| PF3D7_70712400 | Extracellular | FSVYGIVH<br>RLKAVIP<br>IVVGPAYG |

## Discussion

Our novel hybrid deep learning approach, HYDRA, presents a significant advancement in target-aware peptide design, merging the strengths of diffusion models and binding affinity optimization. This hybrid deep learning framework addresses the limitations of existing methods by incorporating a unique iterative optimization algorithm for maximizing binding affinity within an SE(3)-equivariant diffusion generation process. This strategy successfully generates stable peptide binders with high affinity towards target receptors, demonstrating its potential for overcoming challenges associated with peptide therapeutics.

One major hurdle for peptide drugs is their susceptibility to protease degradation, leading to reduced efficacy and short half-life. HYDRA tackles this issue by prioritizing stable interactions between the generated peptides and target receptors during the design process. The model optimizes the binding affinity of the protein-peptide complex while considering structural factors, aiming to generate peptides with enhanced stability and longer half-lives, potentially prolonging their therapeutic effect. Another crucial design consideration is cell permeability, as it directly affects the ability of peptides to reach their target intracellular sites. We address this by incorporating structural properties relevant to cell penetration into the design process. HYDRA generates peptides with a balanced ratio of hydrophilic and hydrophobic residues, enhancing their membrane contact and facilitating efficient cellular uptake. HYDRA goes beyond existing data-driven peptide design approaches by introducing a novel hybrid model that leverages a denoising diffusion model in conjunction with a binding affinity maximization algorithm. This approach utilizes a 3D-structure-specific representation of the protein binding site and the peptide, allowing for the generation of diverse, high-quality peptides tailored to specific binding locations.

To showcase the model’s capabilities, we successfully applied it to design *de novo* peptide binders for proteins expressed by PfEMP1 genes, key contributors to antigenic variation in the malaria parasite. By analyzing Plasmodium parasite single-cell transcriptome data, we identified highly expressed PfEMP1 genes and subsequently pinpointed strong and medium binding sites within their structures using CAVITY^38^. Following peptide generation using HYDRA, we selected a subset of peptides exhibiting superior characteristics for each protein, demonstrating the model’s ability to generate highly stable, cell-permeable, and potentially effective drug candidates.

Overall, our findings demonstrate the remarkable potential of hybrid deep learning approaches like HYDRA in revolutionizing peptide drug discovery. This work paves the way for the development of novel therapeutic peptides with improved stability, efficacy, and targeted delivery, ultimately contributing to advancements in healthcare. In the future, it might be worth exploring the possibility of using fully differentiable local peptide structure optimization and binding affinity computation algorithms^41^ in order to construct a faster, end-to-end deep learning framework for *de novo* design of stable, therapeutic peptides.

## Methods

### Problem Definition

Given a receptor protein and the coordinates of its binding pocket, we aim to computationally design novel, stable peptide binders exhibiting high affinity towards this pocket. We represent the binding pocket as a set of atoms, 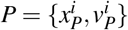, where 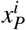 denotes the three-dimensional Cartesian coordinates of each atom in the protein, and 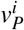 signifies its corresponding feature vector. The chosen feature vector, 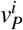, specifically encodes the atom type and its associated amino acid residue. Following this, we aim to generate peptides represented as 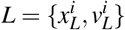. Here, 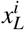 corresponds to the 3D Cartesian coordinates of each atom within the peptide, and *v*^*i*^ signifies its feature vector, similarly encoding atom type and the corresponding amino acid residue.

### HYDRA

To promote the generation of chemically stable peptides for the target receptor, HYDRA employs a two-stage process: (1) amino acid residue generation using a diffusion model (Figure 8) and (2) peptide reconstruction with binding affinity optimization (Figure 9). This necessitates an intermediate representation for the generated residues following the first stage. We represent this intermediate state as 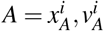, where 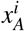 denotes the center-of-mass for each amino acid in the putative peptide and 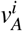 encodes the corresponding amino acid type. Subsequently, the second stage utilizes this intermediate representation as input to reconstruct the final peptide, aiming to maximize its predicted binding affinity towards the target receptor.

**Figure 4.**
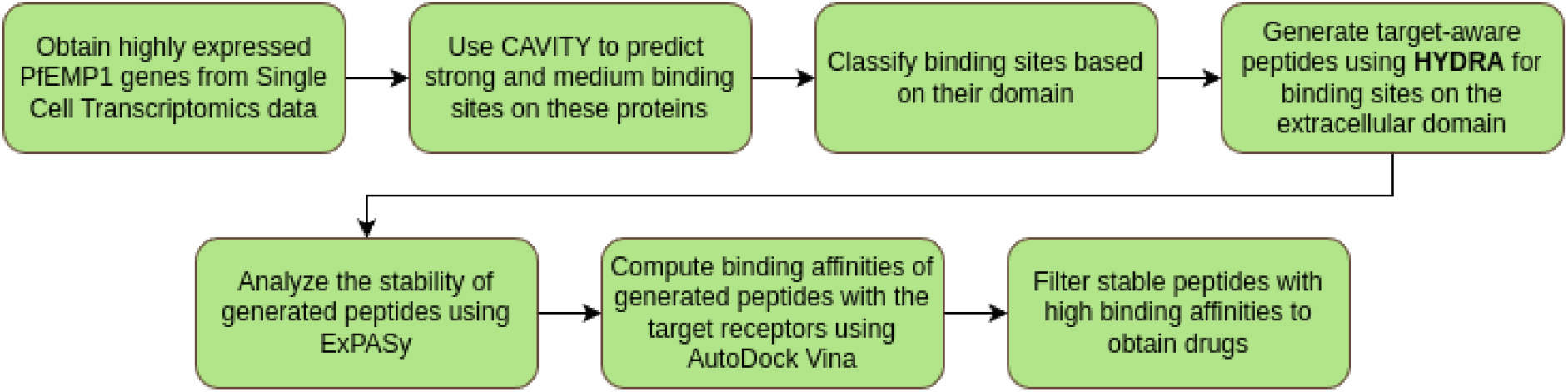
Designing peptides for PfEMP1 receptors. A consolidated flowchart illustrating the binding site detection and *de novo* peptide binder generation pipeline for PfEMP1 Targets

**Figure 5.**
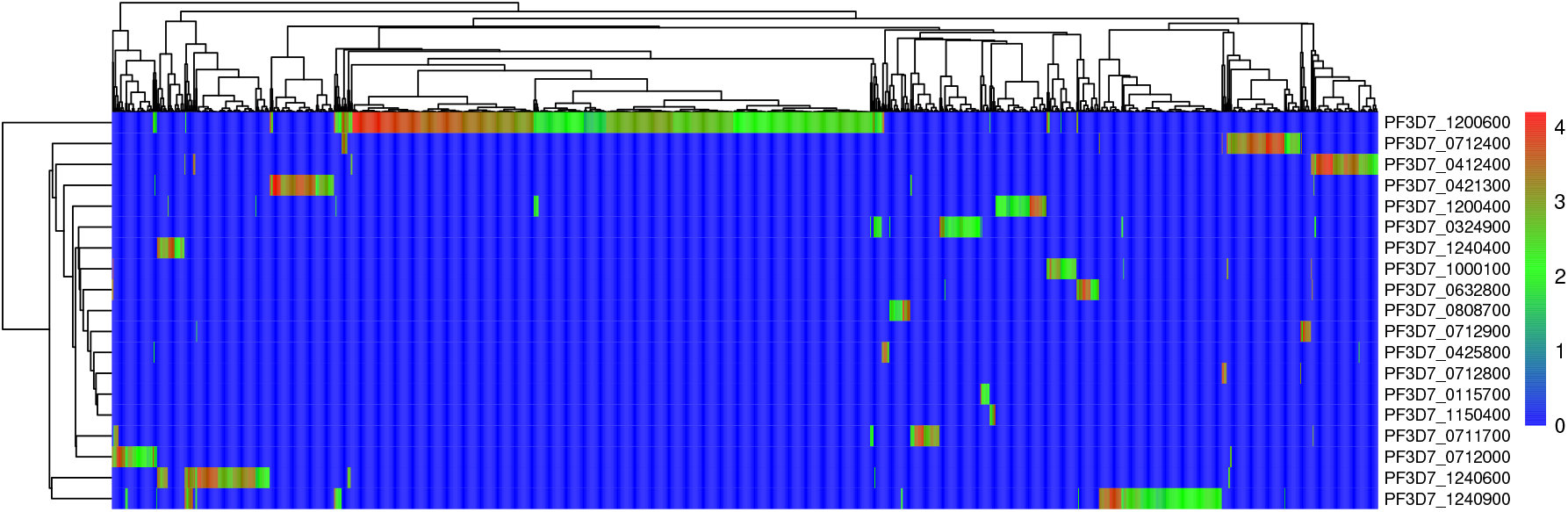
Gene expression profiling of the PfEMP1 family. The proteins PF3D7_1200600, PF3D7_1150400, and PF3D7_70712400 are highly expressed, as observed in the heatmap.

#### Residue Generation

Deep generative modeling has recently surged in popularity due to its ability to generate novel, high-fidelity data across various domains, from drug discovery to artistic creation. Among deep generative models, diffusion models are gaining prominence due to their ability to effectively generate realistic data through a denoising process. Diffusion models are inspired by non-equilibrium thermodynamics^42,43^ and employ a sequential process of noise injection and denoising to learn the underlying distribution of data. During training, the model progressively adds Gaussian noise to real data points, gradually transforming them into isotropic Gaussian noise. This process is modeled as a Markov chain with T discrete steps. Leveraging the Markovian property, the model can efficiently compute the probability density at any given step *t* solely based on the probability density at the preceding step *t* − 1.

Diffusion models have emerged as a promising approach in drug design due to their ability to generate diverse and high-quality 3D molecular structures in a non-autoregressive fashion^1^ as an improvement over sequence-based molecule generation models that make use of Simplified Molecular-Input Line-Entry System (SMILES)^44^ representations that lack detailed spatial information. Inspired by the success of diffusion models in creating 3D molecular structures, we explored their potential in generating peptide shapes specifically designed to fit into target binding pockets. For conciseness, we represent a peptide as a set of amino acid residues *A* = [*x, v*], where [·, ·] is the concatenation operator, *x* ∈ ℝ^*R*×3^ denotes the 3D Cartesian coordinates of the center-of-mass of each amino acid residue, and *v* ∈ ℝ^*R*×*K*^ denotes the one-hot encoded amino acid residue type.

We use a Gaussian distribution (𝒩) for the continuous coordinates and a categorical distribution (𝒞) for the discrete residue types represented as one-hot vectors. The residue distribution is modeled as a product of these individual distributions. A small Gaussian noise and a uniform noise across all categories are added to the coordinates and types, respectively, at each time step *t*:

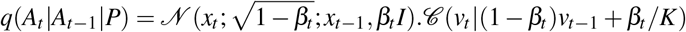

Here, *β*_*i*_ denotes fixed variance schedules, which may differ in practice but are denoted with the same symbol for maintaining conciseness. By employing the reparameterization trick and taking *α*_*t*_ = 1 −*β*_*t*_ and 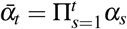, the iterative noise injection process can be significantly accelerated. *A*_*t*_ can now be represented as:

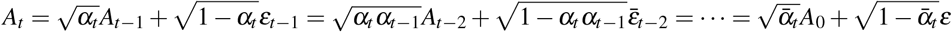

Here, *ε*_*t*−1_, *ε*_*t*−2_, … are noise from 𝒩 (0, *I*) and *ε* combines the noise terms. Consequently

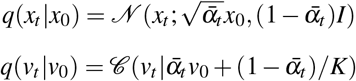

In the denoising process, we aim to recover the initial peptide, *A*_0_, from the final noisy state, *A*_*T*_. However, directly calculating the exact reverse distribution added with the noise is intractable. To address this, we employ a neural network parameterized by *θ* to approximate this reverse distribution.

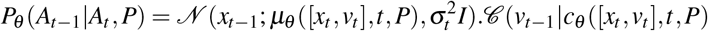

Crucially, this generative process must be invariant to rotations and translations of the protein-peptide complex. This inductive bias ensures consistent likelihood predictions *P*_θ_ (*A*_0_|*P*), essential for accurate 3D peptide structure generation. As informed by existing literature^45,46^, the Markov transition *P*_θ_ (*A*_*t*−1_|*A*_*t*_, *P*_*b*_) must be SE(3)-equivariant and the initial density of our generative process, *P*(*A*_*T*_| *P*_*b*_), is SE(3)-invariant. We need to take into account the amino acid residue coordinates because, during the generative process, atom types are always invariant to SE(3)-transformation. Here, we model [*x*_0_, *v*_0_] to get *µ*_θ_ ([*x*_*t*_, *v*_*t*_], *t, P*) and *c*_θ_ ([*x*_*t*_, *v*_*t*_], *t, P*).

At the *l*-th layer, the hidden embedding **h** and coordinates **x** of the amino acid residues are updated alternately as follows:

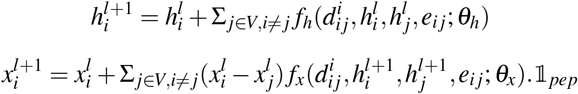

Here, *d*_*i j*_ is the Euclidean distance between residues *i* and *j*, and *e*_*i j*_ denotes a connection between these residues. We use a mask 𝟙_*pep*_ in order to refrain from updating protein atom coordinates. The initial residue embedding **h**^0^ is obtained from an embedding layer that encodes the amino acid information, and the final residue embedding **h**^L^ is used to obtain the final peptide features through a Multi-Layer Perceptron and a Softmax function.

We train the model by minimizing the variational bound on the negative log-likelihood. Due to the Gaussian nature of *q*(*x*_*t*−1_|*x*_*t*_, *x*_0_) and *P*_θ_ (*x*_*t*−1_|*x*_*t*_), the KL-divergence for the coordinate loss admits the closed-form expression:

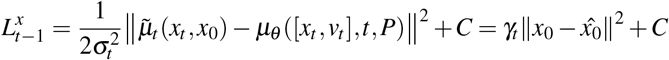

Here, 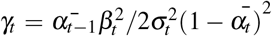 and C is a constant. In practice, training the model with *γ*_t_ = 1 could achieve better performance^43^. We compute the residue type loss directly using the KL-divergence for categorical distributions, given by:

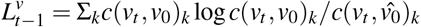

The overall loss function is formulated as a weighted combination of the residue coordinate loss and the residue type loss:

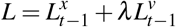

#### Peptide Reconstruction

Following the generation of individual amino acid residues, the next step involves assembling them in the correct sequential order to form complete peptides. To achieve chemically stable protein-peptide complexes, we prioritize the identification of optimal residue connectivity patterns within the generated peptides. This optimization process maximizes the binding affinity of the connected peptide with the target receptor, thereby promoting complex stability. To achieve this, we first compute the distance matrix **D**_*n*×*n*_ such that

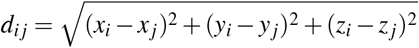

Here, *n* denotes the number of amino acid residues generated that form the peptide, and *d*_*i j*_ denotes each element of the distance matrix that represents Euclidean distance between the centers of mass of residue *i* and residue *j* in 3D space. Due to the symmetry of the distance matrix, the lower triangle can be ignored, leaving 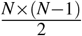 possible edges in the solution space. Following the initial edge identification, a filtering step is applied to eliminate connections exceeding a biologically relevant distance threshold. This step prioritizes the generation of peptides with realistic conformations, as the chemical nature of peptide bonds restricts the maximum distance attainable between adjacent amino acids. Details regarding the threshold calculation can be found in the Supplementary Information. The solution space is reduced to M possible edges following the distance-based thresholding. However, selecting N-1 edges from this set to define the final peptide structure remains challenging. Each edge specifies a connection between residues *i* and *j*, but its directionality determines which residue harbors the C-terminus and, consequently, which residue holds the N-terminus upon bonding.

Construction of a candidate peptide from a chosen set of edges and their directionalities involves a two-stage process. First, the amino acid residues are virtually placed at their predicted center-of-mass positions, and peptide bonds are formed between residues based on the chosen edges, establishing the initial peptide structure. This initial structure then undergoes energy minimization using a Merck Molecular Force Field (MMFF)^47^ to optimize its geometry and achieve a more relaxed, lower-energy conformation. The resulting structure represents the fully reconstructed candidate peptide. Subsequently, the binding affinity between this peptide and the target receptor is assessed using AutoDock Vina^31^. This software performs a local structure optimization of the peptide within the target binding pocket. The optimization employs the Broyden-Fletcher-Goldfarb-Shanno (BFGS) algorithm^48^, followed by the calculation of binding affinity using the Vina Scoring Function^31^:

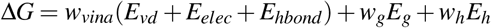

The Vina Scoring Function incorporates the following terms: *E*_vd_, representing the van der Waals interaction energy; *E*_*elec*_, denoting the electrostatic interaction energy; *E*_*hbond*_, accounting for the hydrogen bonding interaction energy; *E*_*g*_, the torsional free energy; and *E*_*h*_, a reference state correction term. These terms are weighted by coefficients *w*_*vina*_, *w*_*g*_, and *w*_*h*_, respectively, which were optimized and assigned during the training of the scoring function^31^.

To identify the peptide with the best binding affinity, an exhaustive evaluation of 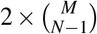 possible amino acid configurations would be required. This becomes computationally intractable as the peptide length (*N*) and, consequently, the solution space (*M* ≥ (*N*− 1)) increases due to the high cost associated with construction, local structure optimization, and binding affinity calculations. This extensive search space necessitates a heuristic-based approach to guide the search towards promising regions.

The peptide reconstruction process inherently lacks a differentiable objective function due to discrete steps in establishing the initial peptide structure, energy minimization using a force field, and local structure optimization by AutoDock Vina. Consequently, gradient-based optimization methods become inapplicable. We opted for a heuristic optimization approach, which treats the objective function as a “black box” and iteratively refines candidate solutions based on their performance within this function. Several no-gradient stochastic optimization algorithms were evaluated, including Genetic Algorithms^49^, Simulated Annealing^50^, and Particle Swarm Optimization (PSO)^51,52^. Binary Particle Swarm Optimization (BPSO) emerged as our preferred choice as we observed faster convergence while achieving comparable results to other methods.

PSO utilizes a population of candidate solutions (particles) representing potential peptide bonds. Each particle has a position in the search space and a velocity guiding its movement. The objective function, which consists of constructing the peptide from the candidate bonds and computing its binding affinity, acts as a fitness function, evaluating each particle’s suitability. Particles track their personal best (*p*_*best*_) position, and the swarm maintains a global best (*g*_*best*_) position found by any particle. Particle velocities are updated iteratively based on their current state, attraction to their *p*_*best*_, and attraction to the *g*_*best*_. This guides them toward promising areas of the search space. Positions are updated based on the new velocity, followed by fitness evaluation. If a particle finds a better position than its *p*_*best*_, it updates its *p*_*best*_. The *g*_*best*_ is updated if a particle discovers a superior position. This cycle repeats for several iterations, allowing the swarm to converge towards optimal solutions iteratively. Binary Particle Swarm Optimization (BPSO) is an implementation of the PSO algorithm where the particles are restricted to the binary domain. A detailed example of the reconstruction workflow for a peptide consisting of 5 amino acid residues is illustrated in Figure 9.

### Model

The SE(3)-equivariant network contains 9 equivariant layers where *f*_*h*_ and *f*_*x*_ are implemented as graph attention layers for features and coordinates with 16 attention heads and 128 hidden features. For Binary PSO, 50 particles are employed with cognitive (c1) and social (c2) acceleration coefficients set to 2.5 and 0.5, respectively. Additionally, the inertia weight is set to 0.9. See Supplementary Materials for details on the diffusion model and PSO optimization.

### Dataset

PepBDB (Peptide Binding DataBase) is a curated structural database specializing in biological peptide-protein interactions^18^. It provides clean data for structure-based peptide drug design, particularly for docking and scoring studies. Compiled from the Protein Data Bank (PDB), PepBDB focuses on structures of interacting peptide-protein complexes, with peptides limited to 50 amino acid residues in length. Regular monthly updates ensure the database reflects the latest data released in the PDB.

### Training

The model was trained using the Adam optimizer^53^, with an initial learning rate of 10^−3^. To prevent overfitting and improve generalization, a data augmentation strategy was employed during training. This involved adding a small Gaussian noise with a standard deviation 0.1 to the protein atom coordinates. Additionally, a learning rate decay schedule was implemented to decay the learning rate exponentially with a factor of 0.6 towards a minimum value of 10^−6^ if there is no improvement in the validation loss for 10 consecutive steps. The model was trained using a batch size of 2, and to balance the contributions of different loss terms within the overall loss function, a factor of *α* = 100 was multiplied onto the residue type loss. Figures concerning the training progression are provided in the Supplementary Materials.

The deep diffusion model was trained on 4x NVIDIA GeForce RTX 2080 Ti GPUs using the Distributed Data Parallel (DDP) Strategy. All inference and reconstruction experiments were carried out on nodes with 40x Intel Xeon E5-2640 v4 CPUs, 80 GB of RAM, and 1x NVIDIA GeForce RTX 2080 Ti GPU belonging to Ada, a High-Performance Computing Cluster at the disposal of the International Institute of Information Technology, Hyderabad, India.

### Peptide generation for PfEMP1 proteins

Plasmodium falciparum employs antigenic variation through switching among its 60 PfEMP1 genes, hindering antimalarial efficacy. We leveraged single-cell RNA-seq data to identify the five most highly expressed PfEMP1 genes. Using CAVITY^38^, we predicted strong and medium druggable binding sites within the proteins encoded by these genes (Figure 6). Subsequently, HYDRA was used to design potential peptide molecules specifically targeted to these binding sites. By analyzing the protein structures, we focused on extracellular and intracellular domains for peptide generation, leveraging cavities as potential binding pockets. Finally, we evaluated the binding affinities between the designed peptides and their target proteins on these PfEMP1 variants.

**Figure 6.**
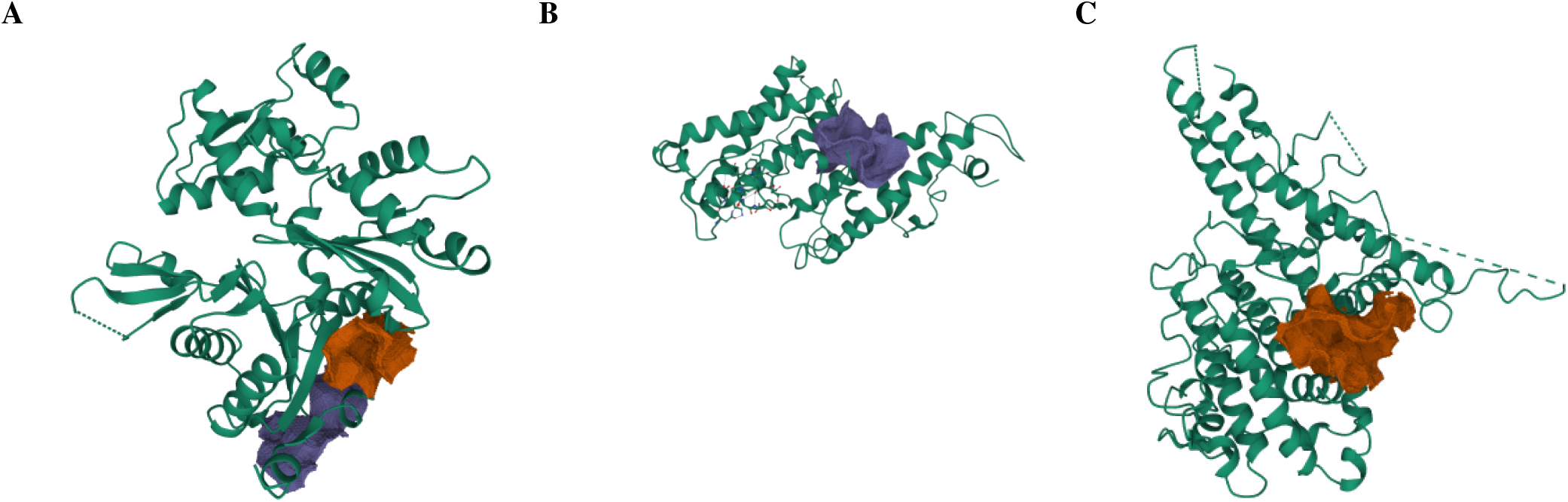
Using CAVITY to identify strong and medium binding sites. Visualization of identified binding sites of (*A*) PF3D7_0712400, (*B*) PF3D7_1200600, and (*C*) PF3D7_1150400. Red and blue regions are binding sites

**Figure 7.**
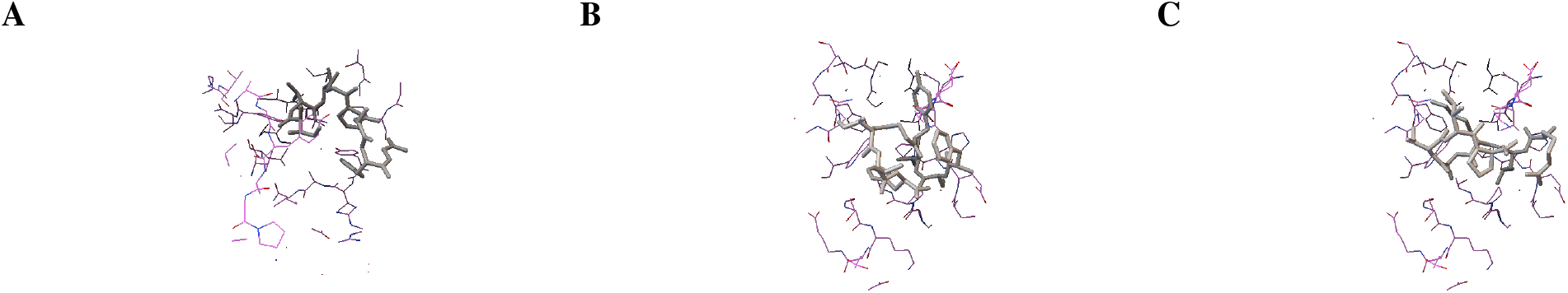
Predicted PfEMP1 protein-peptide complexes. 3D structural representations of the binding sites on PF3D7_0712400 interacting with generated peptides (*A*) FSVYGIVH, (*B*) RLKAVIP, and (*C*) IVVGPAYG.

**Figure 8.**
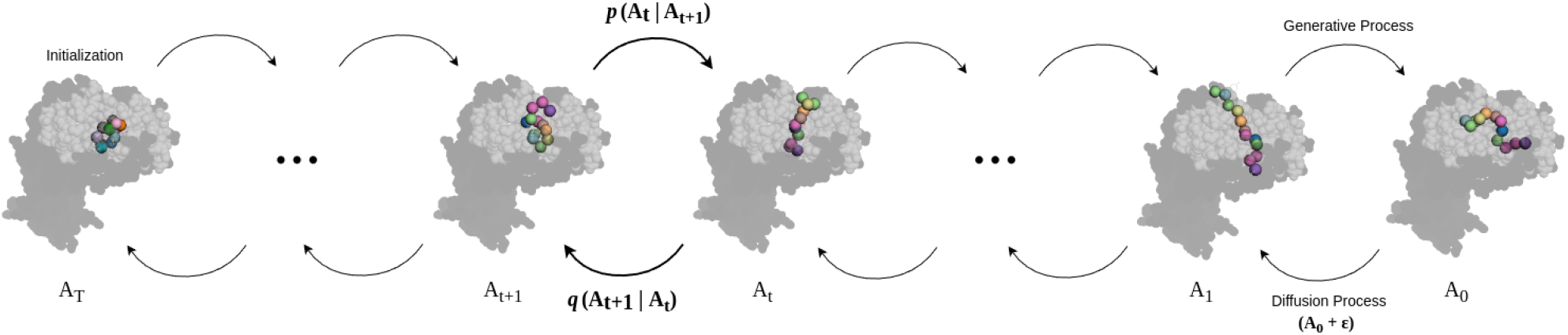
Overview of the diffusion model for residue generation. The diffusion process gradually injects noise into the data, and the generative process learns to recover the data distribution from the noise distribution by means of an SE(3)-equivariant network.

**Figure 9.**
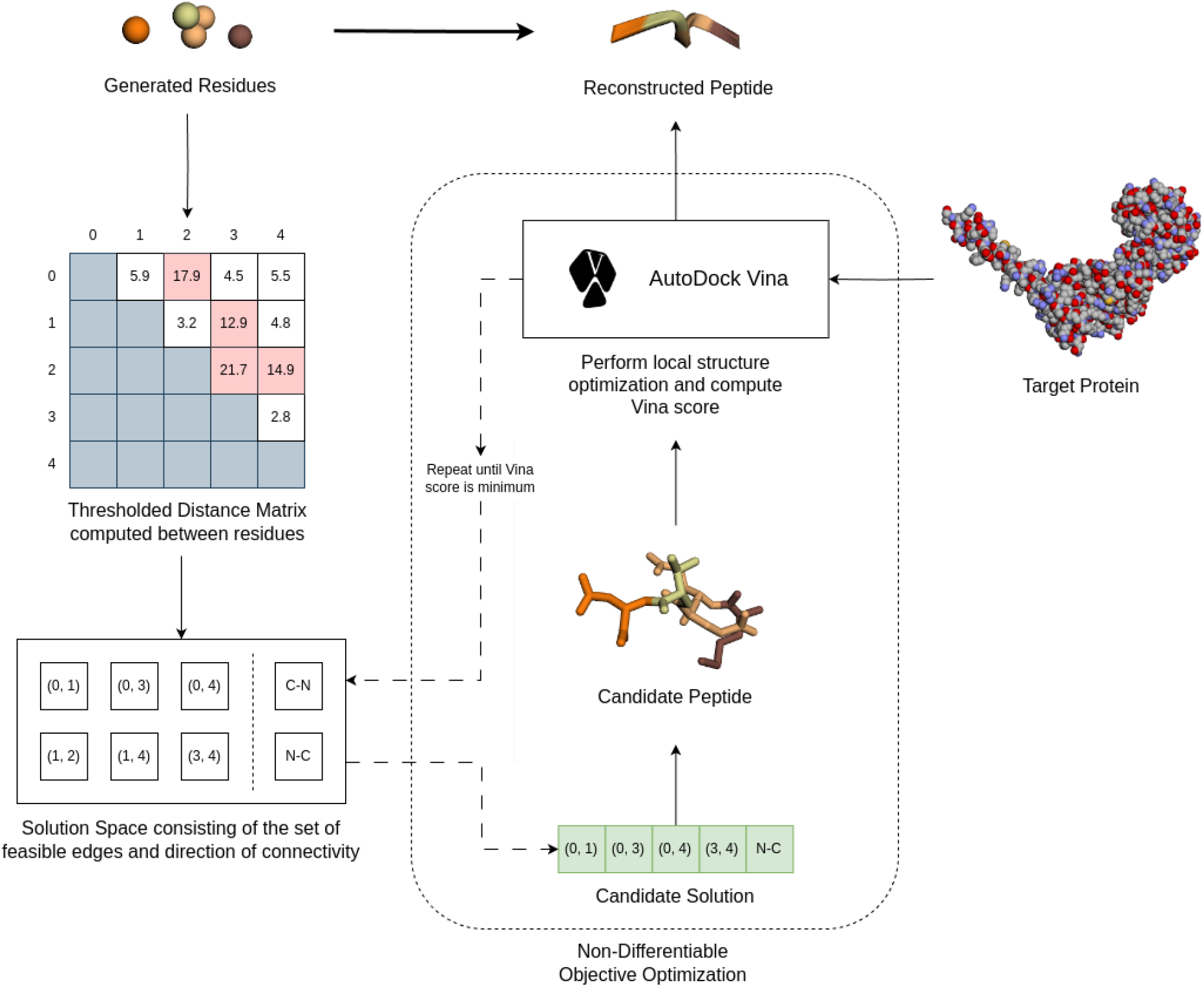
Reconstructing amino acid residue fragments into peptides. First, we calculate distances between amino acid residue centers of mass, forming a distance matrix that captures their spatial relationships. A threshold then filters the matrix, eliminating unrealistic connections and defining feasible peptide conformations. Finally, a non-differentiable optimization algorithm searches this space for the peptide conformation minimizing the Vina score, corresponding to the peptide predicted to form the most stable complex with the receptor.

## Data and Code Availability

PepBDB^18^ served as the foundation for training and testing our diffusion model for protein-peptide interactions. The full dataset can be downloaded from http://huanglab.phys.hust.edu.cn/pepbdb/. Single-cell data to compute highly expressed PfEMP1 genes was collected from VM Howick et al. (2019)^54^. All code for replication and analysis associated with the current submission is available at https://github.com/BhaswarGhoshLab/HYDRA

## Acknowledgements

The authors thank the Department of Biotechnology (No. BT/RLF/Re-entry/32/2017), Government of India, for funding this project.

## Author contributions statement

Conceptualization: S.C., V.S.R & B.G.; methodology: S.C., V.S.R; formal analysis and investigation: V.S.R, S.C. Writing—original draft preparation: S.C., V.S.R; writing—review and editing: S.C., V.S.R & B.G.; funding acquisition: B.G.; supervision: B.G.

## Additional information

**Supplementary Information**: All the code and related supplementary material are available at https://github.com/BhaswarGhoshLab/HYDRA

